# Preconditioning Matters: Enhancing or Impairing Antitumor Immunity with DC Vaccines

**DOI:** 10.1101/2025.05.02.651990

**Authors:** Eric Kwon, Shelby Namen, Colin J. Willoughby, Solomon Kang, Gaurav Pandey, Alexander B. Kim, Carl J. DeSelm

## Abstract

Preconditioning regimens are essential for the success of adoptive cell therapies like CAR T-cells due to effects on the T-cell response, yet they are underexplored and generally absent from cancer vaccine clinical trials. To address this gap, we evaluated the impact of various preconditioning strategies on dendritic cell (DC) vaccine efficacy in a murine tumor model. Mice bearing syngeneic KP tumors expressing ovalbumin received preconditioning with low-dose radiation (LD RT; whole body or tumor only), cyclophosphamide, paclitaxel, LD RT plus cyclophosphamide, or no preconditioning, followed by administration of antigen-loaded DC1s. Tumor growth, survival, and antigen-specific CD8+ T-cell responses were assessed. LD RT preconditioning, whether whole body or tumor-directed, significantly enhanced vaccine-induced antitumor CD8+ T-cell responses and improved survival compared to DC vaccine alone and all other groups. Cyclophosphamide preconditioning reduced vaccine efficacy and negated the benefits of LD RT, while paclitaxel had no significant effect. Notably, whole-body LD RT induced the strongest antigen-specific T-cell response. These findings demonstrate that, similar to CAR T-cell therapy, preconditioning regimens can significantly influence cancer vaccine outcomes. Rational selection of preconditioning agents may either maximize or minimize the therapeutic potential of DC cancer vaccines, and should be considered carefully in clinical trials.

## Introduction

Cancer remains one of the leading causes of morbidity and mortality worldwide, with over 1.6 million new diagnoses in the United States annually^1^. While traditional treatment modalities such as surgery, chemotherapy, and radiation therapy remain mainstays of cancer management, these approaches often lack molecular specificity, lead to significant off-target effects, and may not offer long-term cures, especially in advanced or metastatic cancers. In recent decades, immunotherapy has emerged as a transformative field in oncology, leveraging the immune system’s inherent ability to recognize and destroy malignant cells.

Tumor vaccines—including peptide, mRNA, DNA, and dendritic cell (DC)-based platforms—represent a promising frontier in cancer immunotherapy. Tumor antigen vaccine efficacy depends on endogenous DCs *in vivo* to capture, process, and present the antigen to T cells, thereby initiating an antitumor immune response^2, 3^. In contrast, DC vaccines involve the *ex vivo* generation and maturation of DCs that are directly loaded with tumor antigens prior to administration, effectively bypassing the requirement for antigen uptake *in vivo*, but requiring the ability to culture the appropriate DC population for *in vivo* efficacy. All tumor vaccine strategies ultimately seek to amplify the body’s natural antitumor immunity by providing T cells with precise instructions for recognizing and eliminating cancer cells^4^. Among DC subtypes, conventional type 1 dendritic cells (cDC1s) are uniquely capable of cross-presenting antigens and are essential for the activation of CD8+ cytotoxic T-cell responses against tumors *in vivo*^5^.

A critical, yet underappreciated, aspect of adoptive cell therapies such as CAR T-cell therapy is the use of preconditioning regimens prior to cell infusion. Lymphodepleting agents, most commonly cyclophosphamide, are routinely administered to enhance therapeutic efficacy by creating a favorable immunologic milieu—depleting regulatory and suppressive immune populations, increasing homeostatic cytokine availability, and facilitating the expansion and persistence of transferred T cells^6-8^. Additional agents, such as low-dose radiation^9, 10^ or paclitaxel^11, 12^, have been shown to modulate the tumor microenvironment, increase antigen presentation, and promote T-cell infiltration. The necessity of preconditioning for optimal CAR T-cell efficacy is well established, with improvements in objective response rates and progression-free survival^6-8^.

In contrast, preconditioning regimens are not routinely incorporated into clinical trials of cancer vaccines, despite the mechanistic rationale and potential for similar immunomodulatory benefits. Early-phase clinical studies of DC and other cancer vaccines have demonstrated the capacity to increase antitumor T-cell responses^13-15^, but have not yet translated into clear survival benefits. This discrepancy may, in part, reflect the absence of intentional preconditioning strategies that could potentiate vaccine-induced immunity by overcoming tumor-mediated immune suppression or enhancing T-cell priming and expansion.

To address this critical gap, we systematically evaluated the impact of several preconditioning regimens—including low-dose radiation, cyclophosphamide, and paclitaxel—on the efficacy of tumor antigen loaded DC vaccines in a syngeneic murine tumor model. To minimize confounding variables associated with different vaccine platforms, such as mRNA, DNA, or peptide antigens, as well as the use of various adjuvants, we utilized *ex vivo* antigen-loaded conventional type 1 dendritic cells (DC1s), which are the critical DC population for generating antitumor T-cell responses *in vivo*^5^. This approach allowed us to directly assess the effects of preconditioning on the antigen-induced antitumor immune response, independent of antigen uptake or adjuvant effects. Using ovalbumin-loaded DC1s and established KP tumors, we measured the impact of each preconditioning regimen on tumor growth, survival, and the magnitude of antigen-specific CD8+ T-cell responses. Our objective is to identify clinically translatable preconditioning strategies that can be incorporated into future cancer vaccine trials to maximize antitumor immunity and improve patient outcomes. These findings highlight the importance of rigorously investigating and integrating preconditioning regimens into the design of cancer vaccine therapies.

## Results

### Low-dose radiation as a preconditioning regimen significantly enhances survival of tumor bearing mice

LD RT has proven to be effective in animal studies as a preconditioning regimen for cellular therapies^9, 10^, and we first tested whether the combination of LD RT preconditioning before a DC vaccine is better than simply treating the tumor with LD RT alone, using a tumor model that we cannot get to reject with tumor vaccine alone. The preconditioning regimen was tested on a Kras;Trp53 (KP) pancreatic cancer tumor cells expressing ovalbumin (KP-ova), and a dose of 2 Gy was used for LD RT, since this dose is well below typical clinical treatment doses and is well tolerated by every organ in the human body, but still has cellular and immunomodulatory effects^9, 10^. Four days after subcutaneous tumor injection, LD RT was administered to the mouse. Subsequently, ova-loaded cDC1s were injected intraveneoulsy on day 5, and 8. LD RT alone led to growth reduction but did not fully eliminate any tumors. When LD RT was followed by DC vaccine, all tumors fully rejected, indicated that the addition of DC vaccine to LD RT improves tumor rejection and survival (Fig. 2B). Having established that DC vaccine improves outcomes over LD RT preconditioning alone, we next asked whether LD RT preconditioning improves outcomes over DC vaccine alone, and whether an optimal preconditioning regimen exists among LD RT and two promising chemotherapy agents (cyclophosphamide and paclitaxel) that are also known to modulate the tumor microenvironment.

Cyclophosphamide is a standard preconditioning agent for CAR T-cell immunotherapies^7^, however a major goal of preconditioning for CAR T-cells is to lymphodeplete, leading to space for new T-cells and release of T-cell supporting cytokines which help newly inject CAR T-cells grow and survive. We used a cyclophosphamide dosage of 250 mg/kg, a standard to regimens for other cellular immunotherapies^16^. To see whether these therapies could combine to work more effectively, which is also being tested in clinical trials for CAR T-cells (NCT06623630), the two were both utilized in one mouse cohort. Paclitaxel was utilized in a separate cohort at 20 mg/kg, which has shown promising preconditioning effects in mouse models^12^. Finally, a control group with only DC vaccine administration was included, to test whether preconditioning improves DC vaccine efficacy. All these preconditioning regimens were applied to mice with established tumor and ova-loaded cDC1s were give intravenously on day 5, and 8, at a lower “stress test” dose than in the first experimental design (1e6 and 1.5e5, versus 1e6 and 1e6), to allow for more incomplete responses that better discern subtle differences between the effects of each preconditioning regimen.

Paclitaxel as a preconditioning led to significantly slower tumor growth, but non-significantly improved survival (63 days vs 57 days, respectively) over DC vaccine alone (Fig. 1D). However, the greatest benefit on long-term survival benefit (surviving to day 107) was provided by the LD RT group. Survival in this group was statistically superior to LD RT-cyclophosphamide + DC vaccine, cyclophosphamide + DC vaccine, and DC vaccine only groups. Tumor growth was also significantly reduced with LD RT preconditioning compared with all other preconditioning groups (Fig. 1D).

**Fig. 1:**
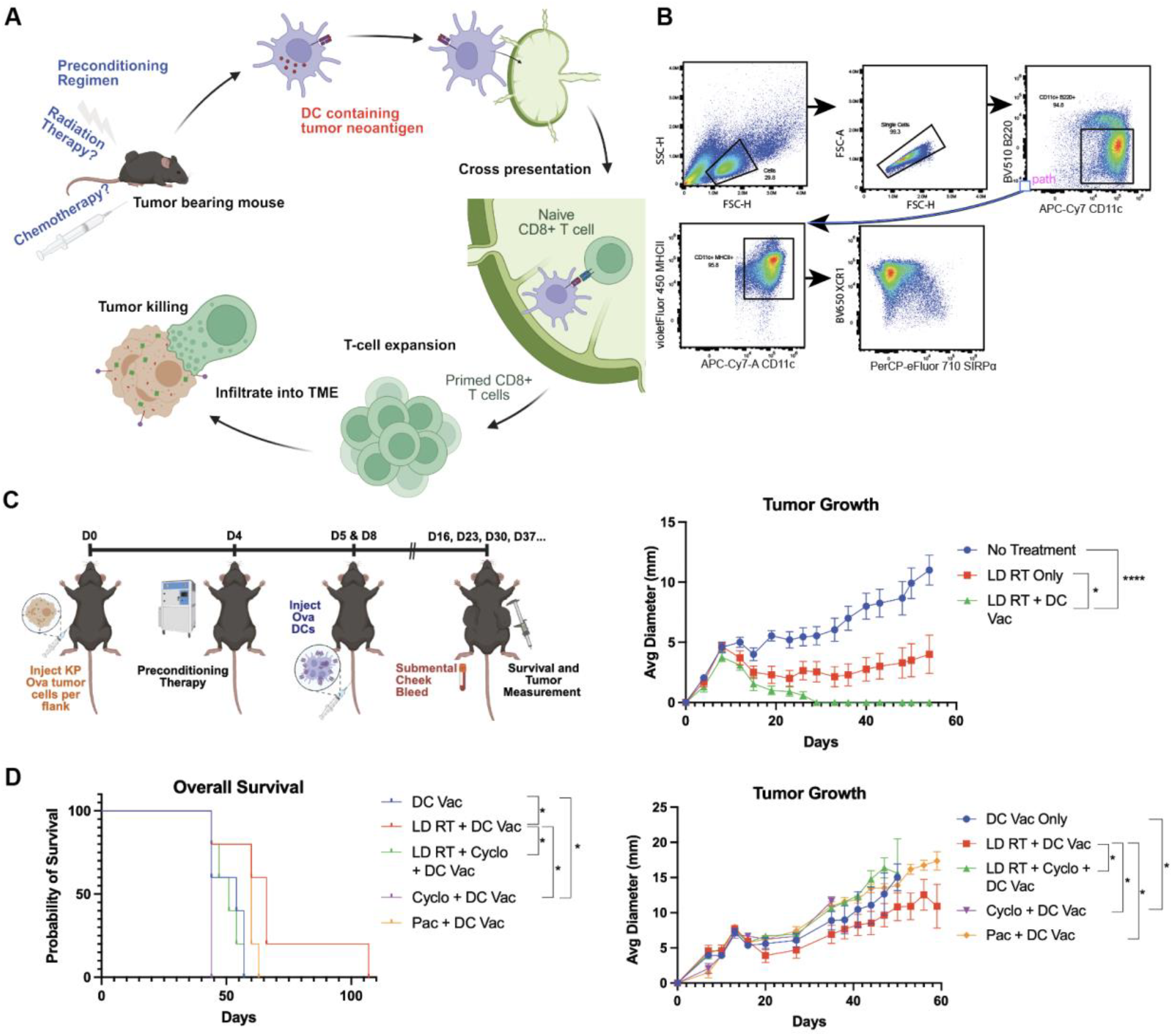
Low-dose radiation as a preconditioning regimen for dendritic cell vaccines significantly improves survival. Mice are injected with ova-expressing KP pancreas tumor cells on day 0. Preconditioning therapy is given on day 4. Bone marrow Flt3-differentiated cDCs are electroporated with the ovalbumin protein (Ova DCs) and injected into tumor bearing mice through the tail vein on day 5 and 8. **A** Mechanistically, the Ova-containing DCs travel to tumor-draining lymph nodes and cross-present the processed SIINFEKL peptide on MHC-I molecules to prime naive CD8 T-cells. Those anti-tumor CD8 T-cells then traffic to tumor sites and perform targeted killing on the Ova-expressing KP Ova tumors. **B** Representative flow cytometry plots of bone marrow differentiated cDCs used in mouse experiments. Sequential gating includes B220^-^ CD11c^+^ MHC II^+^ XCR1^+^ SIRPa^-^ cells. **C** KP Ova cells were injected in C57BL/6 mice on each flank subcutaneously, then irradiated with 2 Gy or not treated. Primary bone marrow cells were isolated from C57BL/6 mice bone marrow, differentiated into cDC1s, and then electroporated with Ova and injected in the tail vein intravenously on day 5 (1e6) and day 8 (1e6). Tumor diameter and mouse survival rates were recorded. Combined survival curves of mice treated with varying preconditioning regimens are shown. Log-rank (Mantel-Cox) tests were conducted to detect significance. **D** KP Ova cells were injected in C57BL/6 mice on each flank subcutaneously, then irradiated with 2 Gy, injected with cyclophosphamide (250 mg/kg), or injected with paclitaxel (20mg/kg). Primary bone marrow cells were isolated from C57BL/6 mice bone marrow, differentiated into cDC1s, and then electroporated with Ova and injected in the tail vein intravenously on day 5 (1e6) and day 8 (1.5e5). Tumor diameter and mouse survival rates were recorded. Log-rank (Mantel-Cox) tests were conducted to detect significant survival differences. Tumor growth was compared using multiple unpaired t-tests with the two-stage step-up method of Benjamini, Krieger and Yekutieli to correct for FDR and multiple comparisons. If tumor diameter was significantly different between groups on two or more occasions, significance was assigned.

### Whole body low-dose radiation and tumor-directed low-dose radiation significantly improve cohort survival

LD RT when utilized with the DC vaccine produced the greatest overall survival of the group as seen in Fig. 1D. LD RT was used to the entire body in these studies for simplicity, but clinically, patients are generally treated with targeted radiation to their sites of known disease. Whether there may be divergent outcomes of using total body LD RT versus tumor-directed LD RT for DC vaccine preconditioning is unexplored. To investigate potential differences from those two modalities of radiation, we treated mice with radiation of 2 Gy to the whole body, to the tumor only, as well as no preconditioning (Fig. 2A). Tumor-bearing mice treated with either total body or tumor only LD RT exhibited similarly reduced rates of tumor growth compared to no preconditioning (Fig. 2B), and both radiation preconditioning approaches significantly improved survival over DC vaccine alone. Compared to DC vaccine alone, whose longest survival was recorded at 62 days, the total body LD RT group included mice that survived throughout the study, surpassing 160 days. Survival outcomes for both radiation modalities, however, were comparable and without statistically significant differences. This is reassuring because it demonstrates that even total body irradiation of 2 Gy does not systemically deplete or damage T-cells enough to hamper the effect of a DC vaccine, and in fact improves the ultimate response.

**Fig. 2:**
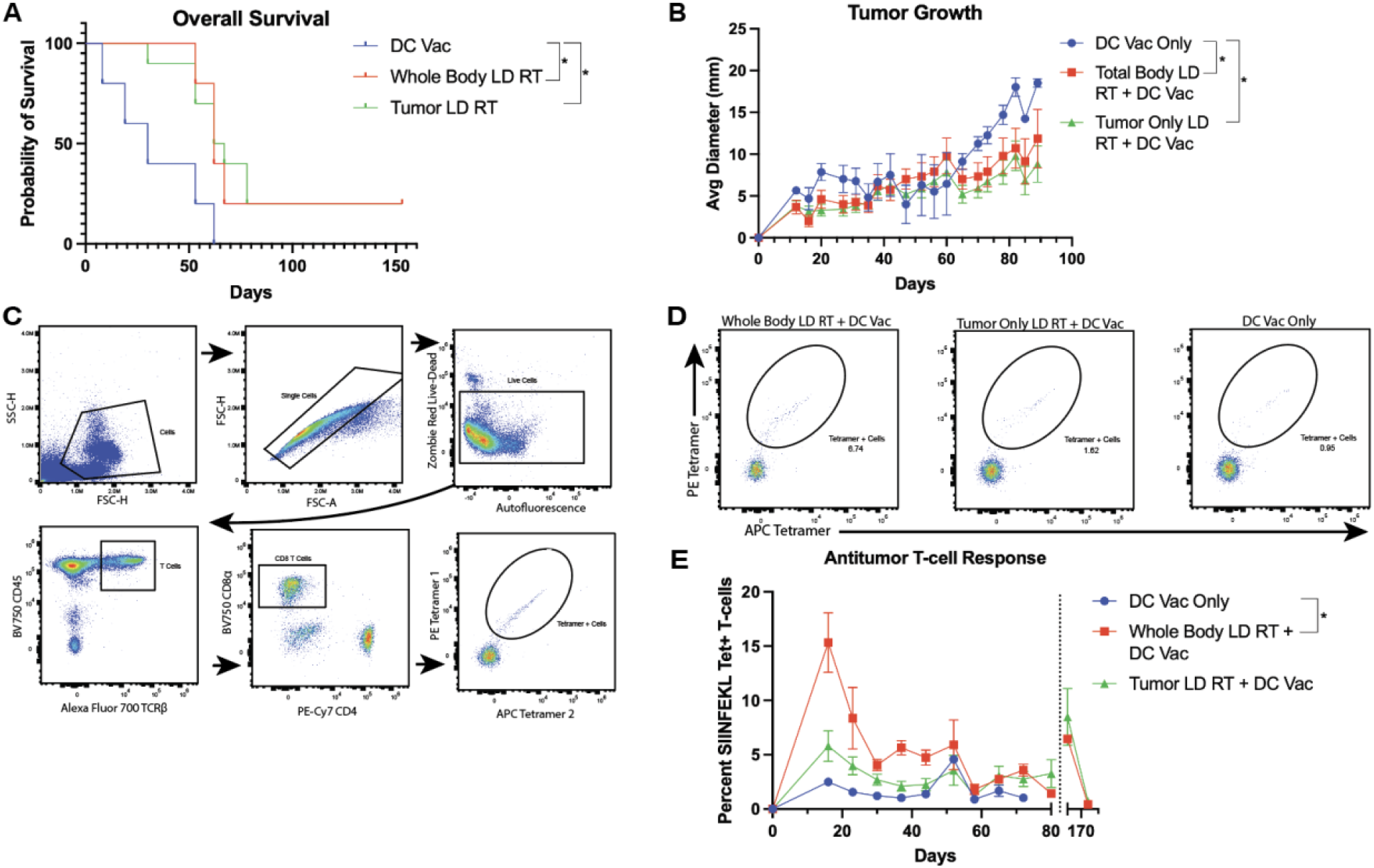
Total body low-dose and tumor-directed low-dose radiation improved responses to DC vaccination and overall survival. **A** KP Ova cells were injected in C57BL/6 mice on each flank subcutaneously, then irradiated with 2 Gy x 1 to the whole body, to the tumor only, or no preconditioning regimen. Primary bone marrow cells were isolated from C57BL/6 mice bone marrow, differentiated into cDC1s, and then electroporated with Ova. Combined survival curves of mice from the same experiment. For comparison of survival curves, the log-rank (Mantel-Cox) test was applied. **B** Tumor diameter recorded over time. **C** Submental cheek bleed samples were stained for T-cell receptors (TCR) that bound the SIINFEKL peptide in Ova, revealing tumor-specific T-cells. Representative gating strategy for identifying tetramer-positive CD8+ T cells from submandibular blood using flow cytometry. Sequential gating includes autofluorescence^-^ Zombie Red^-^ CD45^+^ TCRb^+^ CD8^+^ tetramer^+^ T cells, showing the staining to detect antigen (SIINFEKL)-specific T cells. **D** Representative plots of tetramer (SIINFEKL)-specific T cells from each group. **E** Percentages of tetramer-positive CD8+ T cells of all CD8+ T cells from submental blood are shown with SEM over time. Multiple unpaired t-tests were conducted with the two-stage step-up method of Benjamini, Krieger and Yekutieli to correct for FDR and multiple comparisons. **F** Mice that survived were rechallenged to KP Ova tumor after initial rejection. Percentages of tetramer-positive CD8+ T cells of all CD8+ T cells from submental blood are shown with SEM. These mice re-expressed tumor-specific T-cells and rejected the tumor. In B and E, multiple unpaired t-tests were conducted with the two-stage step-up method of Benjamini, Krieger and Yekutieli to correct for FDR and multiple comparisons and significance was assigned if a significant difference existed between groups on two or more occasions.

### Whole-body LD RT results in significant increases in tumor-specific CD8+ T-cell populations

While survival data is perhaps the most important element, since we know RT can directly affect or kill T-cells, even at low doses, we aimed to define the tumor-specific CD8+ T-cell response after either total body or tumor only LD RT preconditioning and DC vaccine therapy compared with DC vaccine therapy alone. The KP-ova tumor cell line utilized presents ova peptide on its surface MHC, as do the ova-loaded DCs in the vaccine, in the form of a SIINFEKL peptide on MHC I. The ova-presenting dendritic cells should activate SIINFEKL-specific CD8+ T cells systemically, which then target the tumors and slow the growth of the tumor. These T-cells can be recognized by SIINFEKL tetramer, and these tetramer positive T-cells should be an objective measure of the antitumor immune response elicited by IV injected DC vaccine. Submental cheek bleeds were done weekly to analyze the SIINFEKL-specific CD8+ T cells through tetramer staining (Fig. 2C). Whole body LD RT elicited the highest percentage of SIINFEKL-specific CD8+ T-cells amongst all CD8+ T-cells as seen in Fig. 2D-E, with tumor only LD RT prior to DC vaccine eliciting an intermediate tetramer+ T-cell response. While there were no long-term survivors after DC vaccine alone, there were long-term survivors after tumor only and whole-body LD RT preconditioning plus DC vaccine conditions, and to assess for a memory T-cell response these surviving mice were re-challenged to tumor over 150 days later. A re-emergence of SIINFEKL-specific CD8+ T-cells was seen in both the tumor only and total body LD RT preconditioning groups, and re-injected tumor rejected, indicated that antitumor memory T-cells were successfully formed by these therapeutic combinations (Fig. 2F).

## Discussion

We investigated whether preconditioning with cyclophosphamide, paclitaxel, low-dose radiation (LD RT), or LD RT plus cyclophosphamide could enhance DC vaccination against KP-ova tumors. We also compared targeted versus total body LD RT, as the latter may impact immune progenitors. LD RT, whether local or systemic, was more effective at slowing tumor growth and extending survival than DC vaccine alone or with cyclophosphamide or paclitaxel. These results align with prior studies showing LD RT benefits in DC therapy, likely through increased MHC I expression on tumor cells^17^, transient lymphodepletion and homeostatic cytokine release^18, 19^, which together enhance T cell responses.

Cyclophosphamide and paclitaxel did not improve outcomes over DC vaccine alone, though optimal dosing was not explored. Notably, adding cyclophosphamide to LD RT negated the benefit of LD RT, and using cyclophosphamide before DC vaccine was the only condition that led to faster tumor growth as well as shorter survival than vaccine alone, suggesting that while cyclophosphamide is useful in CAR T-cell regimens, it may be detrimental for vaccines relying on endogenous T cells due to its strong lymphodepleting effects.

Both whole-body and tumor-directed LD RT significantly delayed tumor progression and improved survival compared to no preconditioning. Local and systemic LD RT were similarly effective for tumor control, indicating that local modulation may suffice for patients with localized or oligometastatic disease. Systemic (whole body) LD RT, however, induced higher frequencies of circulating antigen-specific (tetramer-positive) CD8+ T cells, possibly due to greater cytokine release or enhanced APC trafficking^17^, though this difference did not translate into better tumor control or survival. This suggests that both approaches may surpass the threshold needed for effective antitumor immunity, and that T cell quality and localization are also important.

Mice cured with LD RT and DC vaccine rejected tumor rechallenge and showed renewed expansion of antigen-specific CD8+ T cells and elimination of the re-injected tumor, indicating durable memory. This demonstrates that LD RT preconditioning with DC vaccine is an effective strategy for short term responses as well as long term surveillance. Previous findings also demonstrate that radiation promotes memory T cell formation through inflammatory mediators like type I interferons and IL-12, which aid in memory precursor differentiation and persistence^20, 21^.

We used OVA-loaded DCs as a controlled model to assess how preconditioning shapes T-cell responses. This approach allows precise evaluation of preconditioning effects and is likely relevant to other antigen delivery strategies, such as DNA, mRNA or peptide vaccines, which also depend on DC-mediated T cell priming. Thus, our findings may inform optimization of a broad range of vaccine platforms.

Overall, our results suggest that localized LD RT is sufficient to render the tumor or tumor microenvironment more amenable to antitumor T-cell responses induced by DC vaccination, while systemic whole-body LD RT may further amplify primed T cell expansion systemically. The lack of additional survival benefit with systemic irradiation, despite higher tetramer-positive T cell numbers, highlights the need to assess T cell quality and localization, not just quantity. These insights can guide clinical strategies to balance efficacy and toxicity by combining systemic or spatially confined preconditioning with DC vaccination for optimal systemic immune activation.

## Methods

### Cell culture

Primary monocytes were isolated from mouse bone marrow. The femurs and tibias of the mice were isolated, then centrifugation was used to isolate the bone marrow. On day 0 of the cell culture, the isolated bone marrow cells were plated into a 10 cm plate in 1.5 × 10^6^ cells/mL using IMDM with HEPES and L-glutamine (Cat. 12440061, Gibco) supplemented with 10% heat inactivated fetal bovine serum (Cat. 10082147, Gibco), 0.1% 2-mercapoethanol (Cat. 21985-023, Gibco), 1% Penicillin-Streptomycin (Cat. 15140122, Gibco), 1% sodium pyruvate (Cat. MT25000CI, Corning), 1% Non-essential amino acids (Cat. MT25025CI, Corning), 2.5 ng/ml rec mouse GM-CSF (Cat. 315-03, Peprotech), and 0.1% S17 FLT3L (Cat. 14–8001, eBioscience). On day 5, 50% of the original volume of IMDM media from day 0 was freshly made without GM-CSF and S17 FLT3L was added to the culture on top of the original media. On day 9, a full media change was conducted with fresh IMDM media including GM-CSF and S17 FLT3L into 0.3 × 10^6^ cells/mL into a T175 flask.

On day 16, a 500 uL sample was taken from the culture to analyze the proportion of cDC1s in the culture. Cells were stained with XCR1 Brilliant Violet 650 (Cat. 148220, BioLegend), MHCII violetFluor 450 (Cat. 20-5321-U100, BD Biosciences), B220 Brilliant Violet 510 (Cat. 583103, BioLegend), SIRPα PerCP-eFluor 710 (Cat. 46-1721-82, Invitrogen), CD11c Brilliant Violet 785 (Cat. 117335, BioLegend)

On day 17, the cells are collected from the culture and electroporated in an Invitrogen Neon NxT with ovalbumin peptide SIINFEKL. After letting the cells rest for 2-3 hours, flow cytometry was done again on the cells using the stain above to determine the cDC1 counts after the electroporation process. Using these numbers, the cells were resuspended into PBS and were then injected into the mouse tumor models through tail vein injections.

This culture was conducted twice, once to inject on day 5 of the *in-vivo* experiment, and once to inject day 8 of the *in-vivo* experiment.

### Mouse Tumor Model

The Kras;Trp53 (KP) Ova cell line was engineered and generously donated by the David DeNardo lab. The cell line is a lung adenocarcinoma that expresses ovalbumin on its surface. These cells used collagen-coated plates and DMEM/F-12 Ham (Cat. D6421, Millipore Sigma) supplemented with 1% sodium pyruvate (Cat. MT25000CI, Corning), and 1% Penicillin-Streptomycin (Cat. 15140122, Gibco). These cells were then injected into C57BL/6 mice on each flank subcutaneously on day 0 of the *in-vivo* experiment.

The mice were shaved for easier access to the subcutaneous layer and measurement of tumor size in the future. These mice under a protocol approved by the Washington University or MSKCC Institutional Animal Care and Use Committee (Prot. 24-0239), according to all relevant animal use guidelines and ethical regulations.

### Irradiation

Animal irradiation was performed using a small animal irradiator (Precision X-Rad 225) with open jaws in the anterior-posterior (AP) direction to deliver 2.0 Gy to the entire animal.

To administer radiation to the tumor only, mice were anesthetized under isoflurane and covered with lead plates, leaving only the tumor exposed.

### Chemotherapy

Cyclophosphamide (Cat. 13849, Cayman Chemical), and Paclitaxel (Cat. T7191, Millipore Sigma) were prepared on day 4 of the *in-vivo* experiment. Cyclophosphamide was resuspended in PBS and injected at 250 mg/Kg. Paclitaxel was titered to 20 mg/kg [9] in 20% 50:50 Cremophor EL: ethanol, 80% saline. Both drugs were injected intraperitoneally into their respective groups.

### Flow Cytometry

All antibodies were titrated. Fc Receptor Binding Inhibitor Antibody Human (Cat. 14-9161-73, eBioscience) was used to block Fc receptors. All flow cytometry was conducted on a Cytek Northern Lights 3000 Flow Cytometer. Forward scatter and side scatter values were set to 25, 75, respectively. All flow cytometry analysis was performed on FlowJo.

### Submental Vein Blood Collection

Submental vein blood collection was done to obtain blood samples on a regular basis without causing permanent damage to the mice. Submental blood collection was conducted on day 16, and every 7 days after. If the submental vein was not accessible or did not yield enough blood, then the submandibular vein was used. Five drops, or around 250 uL, was collected from each mouse and gently mixed in EDTA tubes. The red blood cells were then lysed with RBC lysis buffer (Cat. A10492-01, Gibco) and quenched with PBS. After centrifugation and aspiration of the supernatant, the samples were resuspended in 150 uL FACS buffer and transferred into a 96-well plate. These cells were stained with two SIINFEKL tetramer stains, one conjugated with PE and the other with APC. After the tetramer stains, the rest of the master mix was added with CD3 Brilliant Violet 480 (Cat. 565642, BD Biosciences), CD4 PE cy7 (Cat. BDB563933, Biolegend), CD8 Brilliant Violet 570 (Cat. 100739, BioLegend), TCR-β Alexa Fluor 700 (Cat. 109224, BioLegend), CD45 PE Cy5 (Cat. 103109, BioLegend), CD44 APC Vio 770 (Cat. 130124708, Miltenyl Biotec), and CD62L Brilliant Violet 510 (Cat. 104441, BioLegend). Flow cytometry was then conducted to assess the various qualities in T-cell specificity and activation while also looking at the SIINFEKL-specific cytotoxic CD8+ T-cells that were formed in response to the dendritic cell vaccine.

### Statistics and data

All experimental data are presented as mean ± s.e.m. No statistical methods were used to predetermine sample size. Survival analyses were performed using Kaplan-Meier Log-rank. Statistical analysis was performed on GraphPad Prism 10 software. Schematic Fig.s were made with BioRender.com. All *in-vivo* studies were performed using 5 mice per group.

## Author contributions

EK, SN, and CJD conceptualized experiments. EK, SK, CW wrote the manuscript. EK, SN, SK, CW performed mouse and in vitro experiments. CJD provided experimental and manuscript feedback.

---

**Figure S1.**
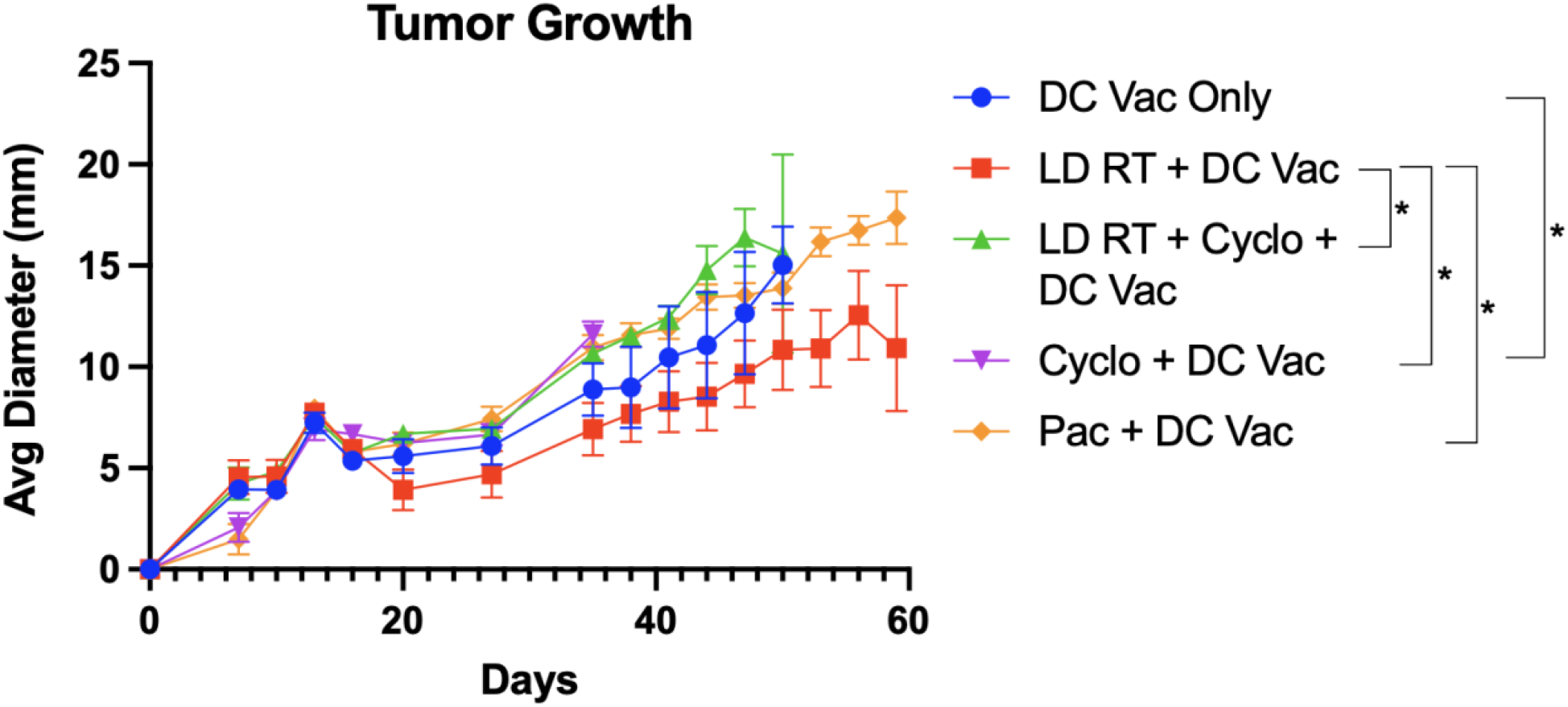
If average tumor diameter was significantly different between two groups on two or more occasions, significance was assigned between groups. Significance was calculated by unpaired *t*-tests, controlled for false discovery rate by two-stage step-up (Benjamini, Krieger, and Yekutieli) to correct for multiple comparisons.

## References

1. Sherman, R.L. et al. Annual Report to the Nation on the Status of Cancer, featuring state-level statistics after the onset of the COVID-19 pandemic. Cancer 131, e35833 (2025).

2. Saadeldin, M.K., Abdel-Aziz, A.K. & Abdellatif, A. Dendritic cell vaccine immunotherapy; the beginning of the end of cancer and COVID-19. A hypothesis. Med Hypotheses 146, 110365 (2021).

3. Zanna, M.Y. et al. Review of Dendritic Cells, Their Role in Clinical Immunology, and Distribution in Various Animal Species. Int J Mol Sci 22 (2021).

4. Palucka, K. & Banchereau, J. Dendritic-cell-based therapeutic cancer vaccines. Immunity 39, 38–48 (2013).

5. Ferris, S.T. et al. cDC1 Vaccines Drive Tumor Rejection by Direct Presentation Independently of Host cDC1. Cancer Immunol Res 10, 920–931 (2022).

6. Turtle, C.J. et al. Immunotherapy of non-Hodgkin’s lymphoma with a defined ratio of CD8+ and CD4+ CD19-specific chimeric antigen receptor-modified T cells. Sci Transl Med 8, 355ra116 (2016).

7. Lickefett, B. et al. Lymphodepletion - an essential but undervalued part of the chimeric antigen receptor T-cell therapy cycle. Front Immunol 14, 1303935 (2023).

8. Ju, A., Choi, S., Jeon, Y. & Kim, K. Lymphodepletion in Chimeric Antigen Receptor T-Cell Therapy for Solid Tumors: A Focus on Brain Tumors. Brain Tumor Res Treat 12, 208–220 (2024).

9. Kim, A.B. et al. Intrinsic tumor resistance to CAR T cells is a dynamic transcriptional state that is exploitable with low-dose radiation. Blood Adv 7, 5396–5408 (2023).

10. DeSelm, C. et al. Low-Dose Radiation Conditioning Enables CAR T Cells to Mitigate Antigen Escape. Mol Ther 26, 2542–2552 (2018).

11. Hsu, F.T., Chen, T.C., Chuang, H.Y., Chang, Y.F. & Hwang, J.J. Enhancement of adoptive T cell transfer with single low dose pretreatment of doxorubicin or paclitaxel in mice. Oncotarget 6, 44134–44150 (2015).

12. Zhong, H. et al. Low-dose paclitaxel prior to intratumoral dendritic cell vaccine modulates intratumoral cytokine network and lung cancer growth. Clin Cancer Res 13, 5455–5462 (2007).

13. Fu, C. & Jiang, A. Dendritic Cells and CD8 T Cell Immunity in Tumor Microenvironment. Front Immunol 9, 3059 (2018).

14. Yu, J., Sun, H., Cao, W., Song, Y. & Jiang, Z. Research progress on dendritic cell vaccines in cancer immunotherapy. Exp Hematol Oncol 11, 3 (2022).

15. Sethna, Z. et al. RNA neoantigen vaccines prime long-lived CD8(+) T cells in pancreatic cancer. Nature 639, 1042–1051 (2025).

16. Khan, M.G.M. & Wang, Y. Advances in the Current Understanding of How Low-Dose Radiation Affects the Cell Cycle. Cells 11 (2022).

17. Burnette, B.C. et al. The efficacy of radiotherapy relies upon induction of type i interferon-dependent innate and adaptive immunity. Cancer Res 71, 2488–2496 (2011).

18. Zebertavage, L.K., Alice, A., Crittenden, M.R. & Gough, M.J. Transcriptional Upregulation of NLRC5 by Radiation Drives STING- and Interferon-Independent MHC-I Expression on Cancer Cells and T Cell Cytotoxicity. Sci Rep 10, 7376 (2020).

19. Lin, C.C. et al. Potentiation of the immunotherapeutic effect of autologous dendritic cells by pretreating hepatocellular carcinoma with low-dose radiation. Clin Invest Med 31, E150–159 (2008).

20. Deng, L. et al. Irradiation and anti-PD-L1 treatment synergistically promote antitumor immunity in mice. J Clin Invest 124, 687–695 (2014).

21. Matsumura, S. et al. Radiation-induced CXCL16 release by breast cancer cells attracts effector T cells. J Immunol 181, 3099–3107 (2008).

